# Domain General Processes for Interactive Touch

**DOI:** 10.1101/2021.09.23.461532

**Authors:** Tao Buck, Courtney DiCocco, Jennifer L. Cuzzocreo, J. Adam Noah, Xian Zhang, Joy Hirsch

## Abstract

The nexus model of social processing proposes that the right temporal parietal junction (rTPJ) serves as a neural hub for cognitive social functions. We test the hypothesis that the rTPJ is a domain general region including somatosensory social functions. Neuroimaging findings and cross-brain coherence for right- and left-hand handclasps with real vs. simulated hands were consistent with the domain general model.

## 1 Introduction

Touch plays a fundamental role in natural social and interactive communication. For example, the handshake is a social act wherein two individuals make contact to greet one another or to conclude a social interaction (Hall & Spencer Hall, 1983). It is commonly used as a proxy for contract agreement that conveys trust (Dolcos et al., 2012). Importantly, a handshake creates an interactive union between two individuals and represents a form of nonverbal social communication (Hall, 2003). Similar to communicating through eye contact or a verbal dialogue, the handshake is an action that has an intention and involves interpersonal feedback. Understanding the neural mechanisms of natural interactions has been challenging due, in part, to technological limitations associated with observations of live interplay between dyads. However, these limitations have been largely resolved by advances in functional near infrared spectroscopy (fNIRS), a non-invasive head mounted spectral absorbance approach, that measures task-related changes in blood oxygen levels in both oxyhemoglobin and deoxyhemoglobin through optical sensors (Jöbsis, 1977; Strangman et al., 2002; Villringer & Chance, 1997). This method allows the recording of hemodynamic signals from two participants simultaneously and has enabled an increased focus on live and natural interactions (Cui et al., 2012; Hasson & Frith, 2016; Hasson et al., 2012; Hirsch et al., 2018; Redcay et al., 2012; Schilbach et al., 2013).

Social touch, such as a handshake or a hand clasp, conveys both somatosensory and interpersonal social information. Although somatosensory pathways are well known, the neural systems associated with social touch information are less well characterized. We distinguish between two alternative models for somatosensory social information: in one case this social information is processed within the contralateral primary and association systems; and in the alternative case the social information is processed separately from the somatosensory information. The Nexus model of social behavior (Carter & Huettel, 2013) proposes that the right temporal parietal junction (rTPJ) integrates multiple streams of cognitive and social information in order to provide a social context, including theory-of-mind functions. Previous studies have found that the right TPJ is also sensitive to live interactions, including eye contact, consistent with social interactive behavior (Kelley et al., 2021; Noah et al., 2020). These visual processes, also with a social meaning, suggest that the rTPJ may be a domain general region for social and interactive processes.

To test this hypothesis for social touch, we use the sensorimotor system and the social task of hand clasping. We employ a hand clasp with a live partner and compare responses with a hand clasp with a simulated hand in order to distinguish a “social” condition from a “non-social” condition. The design of this study takes advantage of the anatomically crossed sensory and motor pathways where expected responses for touch include contralateral regions in sensorimotor hand areas. The hypothesis of a domain general function for rTPJ would be supported if the right TPJ is also be active during social hand-clasping (either right or left hand) and is greater for real hands rather than for simulated hands. In this investigation, fNIRS was used to concurrently record hemodynamic signals from dyads during left and right hand-clasping intended to represent the initial connection made in a common handshake.

## 2 Methods

### 2.1 Participants

Thirty healthy adults (15 pairs; 19 females; mean age: 28.8 ± 12.2 years; 29 right-handed (Oldfield, 1971)), took part in this study. There were 6 female-female, 2 male-male, and 7 female-male pairs. No known neurologic or psychiatric conditions were reported. Written informed consent was provided by all participants in accordance with the guidelines approved by the Yale University Human Investigation Committee (HIC #1501015178), and each person was compensated for participating. Participants who were previously unacquainted were recruited and paired prior to the experiment. All participants had performed a finger-thumb tapping screening task at an earlier date to confirm the presence of recordable hemodynamic signals. See Table S1 for individual participant and group summary demographics.

### 2.2 Paradigm and Procedures

Dyads participated in four tasks (Figure 1A & 1B) during which they clasped right or left hands, as if to initiate a handshake, with either a partner or a simulated hand (i.e., they clasped a real hand or simulated hand with their right hand or left hand). Partners were seated at a table approximately 140 cm from one another. An occluder panel on the table between participants prevented them from seeing each other but allowed them to clasp the other person’s hand in the Real Hand conditions. Simulated hands were arranged in the middle of the table where the partners’ hands met. The order of tasks was arranged randomly, and each condition was repeated twice before proceeding to the next. The time-series of each condition (Figure 1C) is similar to that used in prior studies (Hirsch et al., 2018; Hirsch et al., 2017; Noah et al., 2020; Piva et al., 2017). All runs consisted of 30-second blocks with the active task period alternating with a 15-second rest/baseline period for a total of 180 seconds per condition. An initial auditory cue instructed participants to grasp the simulated hand in front of them or grasp the other participant’s hand. Every three seconds, an auditory tone prompted the dyad to hold or let go of the hand. Participants focused their gaze on a crosshair located on the occluder during the 3 s in which the participants were not clasping hands (real or simulated) and during the 15 s rest/baseline period.

**Figure 1.**
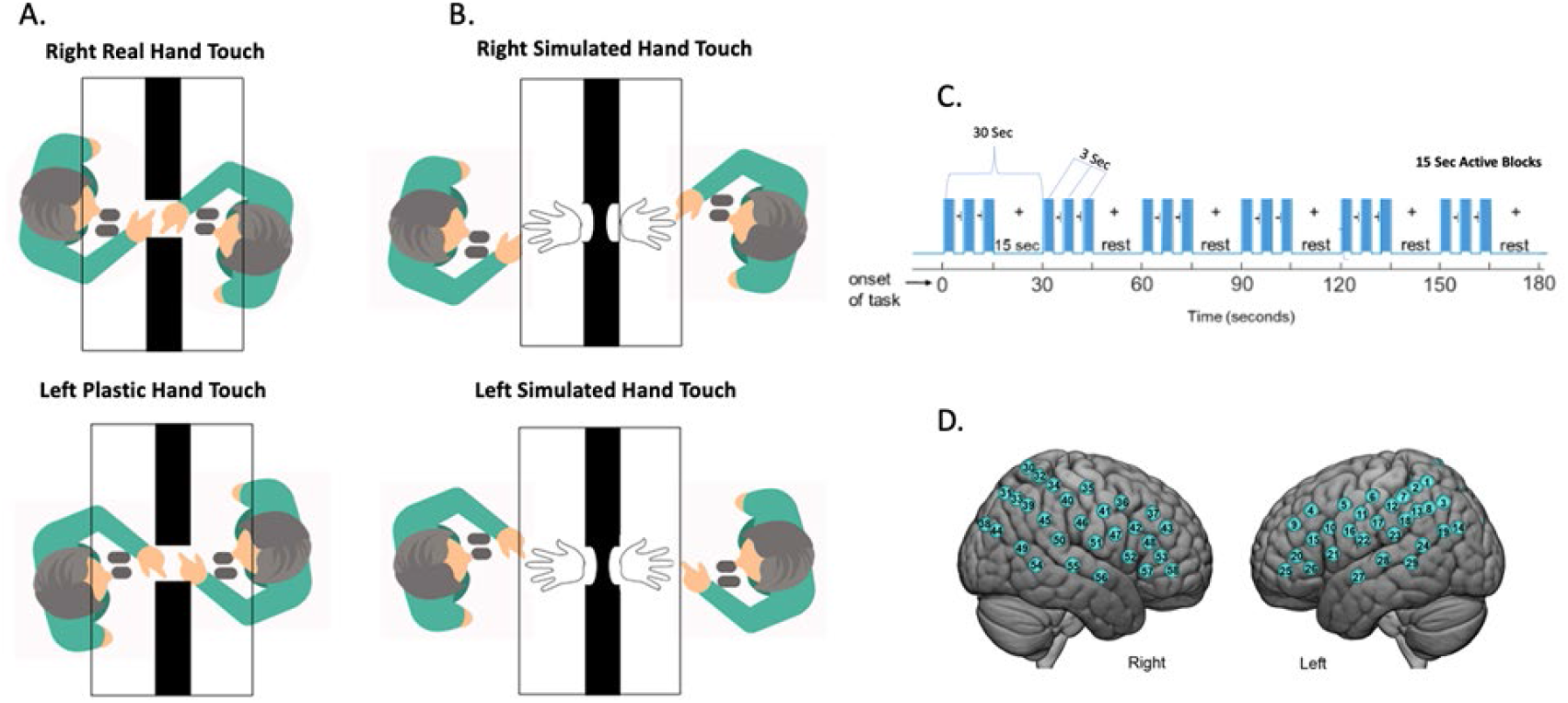
Experimental conditions, time series, and channel layout. **A.** Real Hand condition. Partners faced each other at a distance of 140 cm with an occlusion between them that prohibited them from seeing one another but allowed hand contact. Participants were asked to grasp their partner’s left and right hand. **B.** Simulated Hand condition. Participants were asked to grasp a simulated plastic hand arranged on the table in front of them in the same location as their partner’s hand during the real hand condition. **C.** Time course of the experimental paradigm. The entire duration of every run was three minutes, and each run was repeated twice for the Real Hand and Simulated Hand (right and left hands), for a total of eight runs. Each run included six alternating 15-second task and rest periods. In the task period (blue bars), participants grasped their partner’s hand or a simulated hand in three-second periods alternating with three-second periods of gaze fixation on a crosshair located on the occluder at eye level. During the 15-second rest period, participants looked at a crosshair and rested the hands on the table. **D.** Channel layout. Right and left hemispheres of a single rendered brain illustrate median locations (blue dots) for 58 channels per participant. Montreal Neurological Institute (MNI) coordinates were determined for each channel (numbers on blue dots) by digitizing emitter and detector locations in relation to anterior, posterior, dorsal, and lateral fiduciary markers based on the standard 10-20 system.

### 2.3 Signal Acquisition and Channel Localization

Signal acquisition, optode localization, and signal processing, including global mean removal, were similar to methods described previously (Dravida et al., 2018; Dravida et al., 2020; Hirsch et al., 2018; Hirsch et al., 2017; Kelley et al., 2021; Noah et al., 2017; Noah et al., 2015; Noah et al., 2020; Piva et al., 2017; Zhang et al., 2017; Zhang et al., 2016) and are consistent with recommended standards of practice (Yücel et al., 2021). Hemodynamic signals were acquired using an 80-fiber multichannel, continuous-wave fNIRS system (LABNIRS, Shimadzu Corp., Kyoto, Japan) that emits and detects three wavelengths of light (780 nm, 805 nm, 830 nm). Each participant was fitted with an optode cap with predefined channel distances between holders. Three sizes of caps were used based on the circumference of the heads of subjects. Large caps had a 60 cm circumference. Medium caps were 56.5 cm and small caps were 54.5 cm. Optode distances of 3 cm were designed for the 60 cm cap layout but were scaled equally to smaller caps. To remove hair from each channel opening, a lighted fiber-optic probe (Daiso, Hiroshima, Japan) was used prior to optode placement. Forty emitter and detector optode pairs were organized to record from 58 channels per participant (Figure 1D). Using a modified Beer-Lambert equation (Hazeki & Tamura, 1988; Hoshi, 2003; Matcher et al., 1995), the absorbed light measured by the detectors was translated into concentrations of oxyhemoglobin (OxyHb) and deoxyhemoglobin (deOxyHb). Signals were acquired at 30 Hz. Resistance was evaluated and adjusted for each channel before starting the experiment to ensure that the data obtained would have an acceptable signal-to-noise ratio. Anatomical locations of optodes in relation to standard head landmarks were determined for each participant using a Patriot 3D Digitizer (Polhemus, Colchester, VT) (Eggebrecht et al., 2014; Eggebrecht et al., 2012; Ferradal et al., 2014; Matcher et al., 1995; Okamoto & Dan, 2005; Singh et al., 2005). Montreal Neurological Institute (MNI) coordinates (Mazziotta et al., 2001) for each channel were obtained using NIRS-SPM software (Ye et al., 2009). Anatomical regions corresponding to channel locations were identified with WFU Pickatlas (Maldjian et al., 2004; Maldjian et al., 2003).

### 2.4 Signal Processing and statistical analysis

Channels containing excessive noise were determined and rejected by calculating any channel with raw data having a root mean square 10 times larger than the sum of all the channels root mean square. Data were then converted to concentrations of oxyhemoglobin (OxyHb) and deoxyhemoglobin (deOxyHb) chromophore concentrations using the Beer-Lambert Law (Hazeki & Tamura, 1988; Hoshi, 2003; Matcher et al., 1995). The wavelet detrending function available in NIRS-SPM software was applied to remove baseline drift (Ye et al., 2009). A principal component analysis (PCA) and global-mean filter (Zhang et al., 2017; Zhang et al., 2016) was applied to remove global components due to physiologic factors including blood pressure and other non-neural components (Tachtsidis & Scholkmann, 2016) prior to a general linear model (GLM) analysis. No additional pre-processing steps were performed.

General linear modeling analysis was performed on the hemoglobin difference signal ([Hbdiff] = [HbO**_2_**]- [HHb]) (Tachtsidis et al., 2009), referred to here as the combined signal, where the signs of both the OxyHb (HbO**_2_)** and deOxyHb (HHb) signals are consistent with true brain activity. Event periods (Figure 1C) were convolved with the canonical hemodynamic response function supplied by SPM8 (Penny et al., 2011) to create modeled time courses for obtaining beta values using the GLM approach. The differences between the individual beta values across conditions were interpolated into an MNI brain template for second level analysis, yielding contrast results. Significance was determined using one-tailed t-tests. Findings were corrected using False Discovery Rate (FDR) at p ≤ 0.05 within MATLAB.

A finger-thumb tapping localizer task was performed to confirm the expected hand/finger localizations (Dravida, et al, 2018). Fifteen sec epochs of finger-thumb tapping alternated with 15 secs of rest for a total of six cycles and 180 sec. Two runs were acquired for each hand. The order of hands was counterbalanced. Observation of the expected contralateral pre-motor and supplementary motor localizations for right- and left-hand activity compared to rest is shown in Figure 2 and Table 1A and 1B.

**Figure 2.**
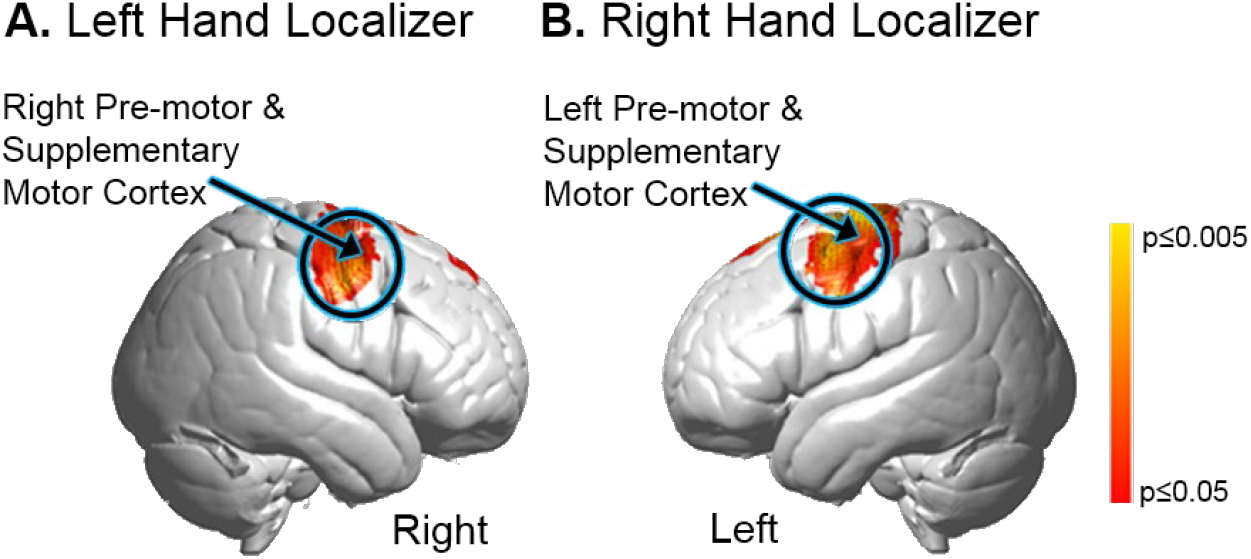
**A.** Right Hand Localizer task: contrast comparison [Real Hand > Rest] based on the combined OxyHb and deOxyHb signals (p≤0.05). Activity was observed in left pre-motor and supplementary motor cortex (See Table 1A). **B.** Left Hand Localizer task: contrast comparison [Real Hand > Rest] based on the combined OxyHb and deOxyHb signals (p≤0.05). Activity was observed in right and left pre-motor and supplementary motor cortex (See Table 1B).

**Table 1A.**
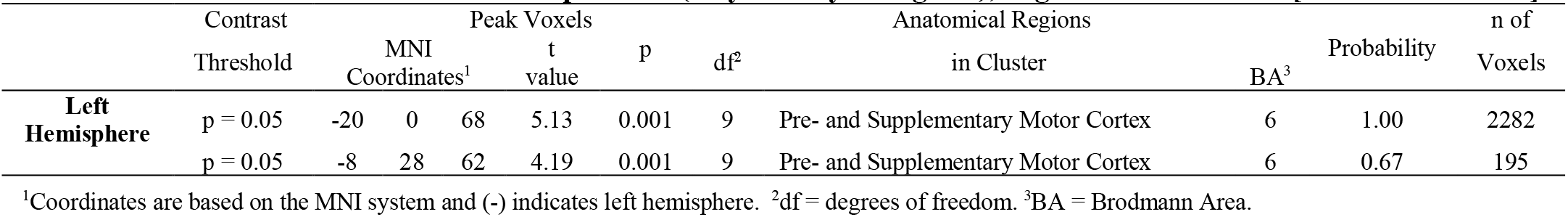
Voxel-wise GLM Contrast comparisons (Oxy+deOxyHb signals), Right Hand Localizer [Real Hand > Rest].

**Table 1B.**
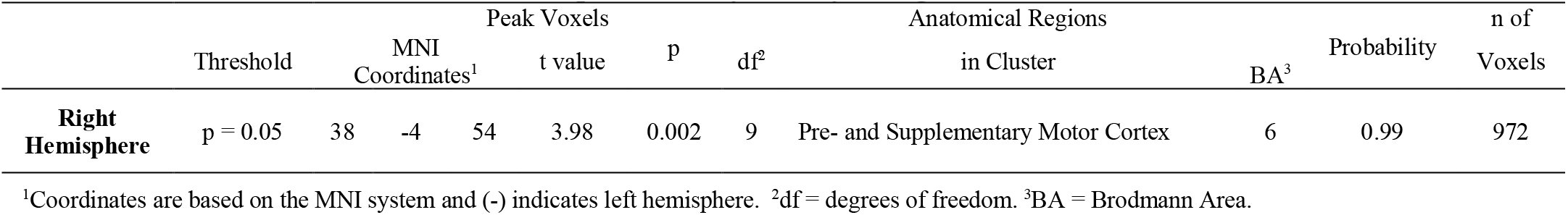
Voxel-wise GLM Contrast comparisons (Oxy+deOxyHb signals), Left Hand Localizer [Real Hand > Rest].

Cross-brain synchrony (coherence) was analysed for the real hand condition versus the simulated hand using wavelet analysis (Torrence & Compo, 1998) from the MATLAB 2018A Wavelet Toolbox. Signals were decomposed into a range of temporal frequencies. Coherence was calculated for each frequency component (y-axis) as a function of component wavelength (period, x-axis) (Hirsch et al., 2018; Hirsch et al., 2021; Noah et al., 2020).

## 3 Results

### 3.1 GLM Analysis

A voxel-wise general linear model (GLM) contrasting the tasks involving the real hand versus the simulated hand combining left and right hands shows activation in the temporoparietal junction in the right hemisphere (Figure 3 and Table 2). These areas include the right superior temporal gyrus (STG), middle temporal gyrus (MTG), and angular gyrus (AG) consistent with social and interactive processing in the rTPJ confirming the domain general hypothesis

**Figure 3.**
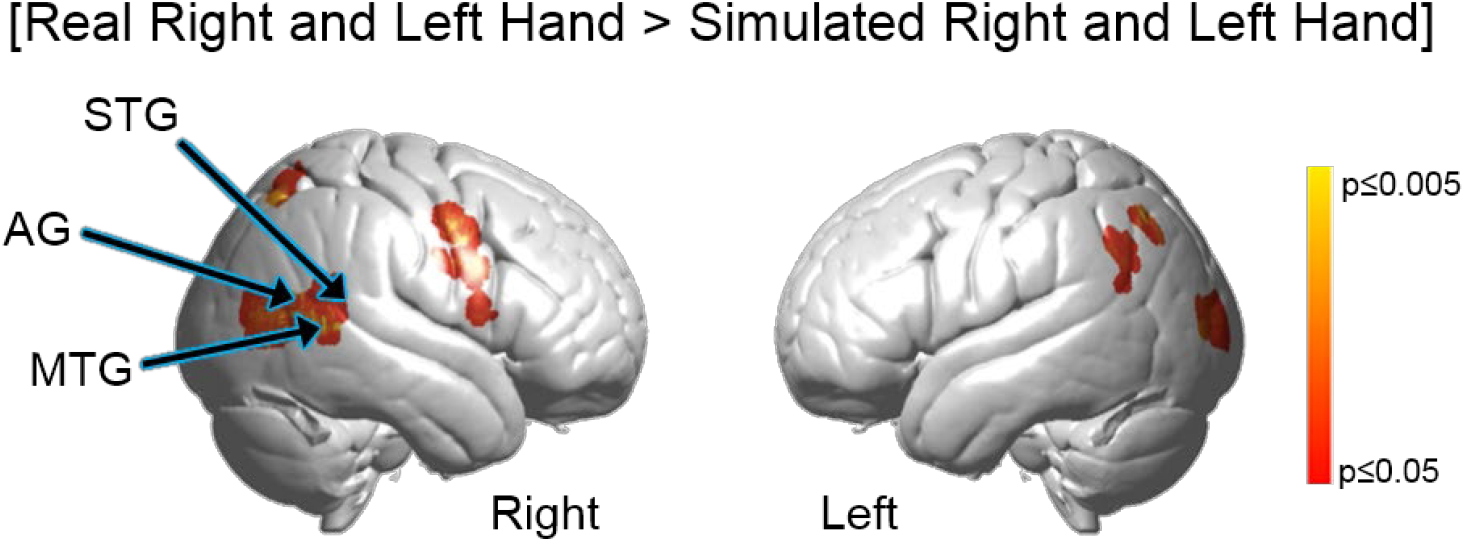
Contrast comparison of all [Real Right and Left Hand > Simulated Right and Left Hand] activity, combining both right and left hands, based on the combined OxyHb and deOxyHb signals (p≤0.05). Activity was observed in right hemisphere superior temporal gyrus (STG); middle temporal gyrus (MTG); and angular gyrus (AG), components of the rTPJ. See Table 2.

**Table 2.**
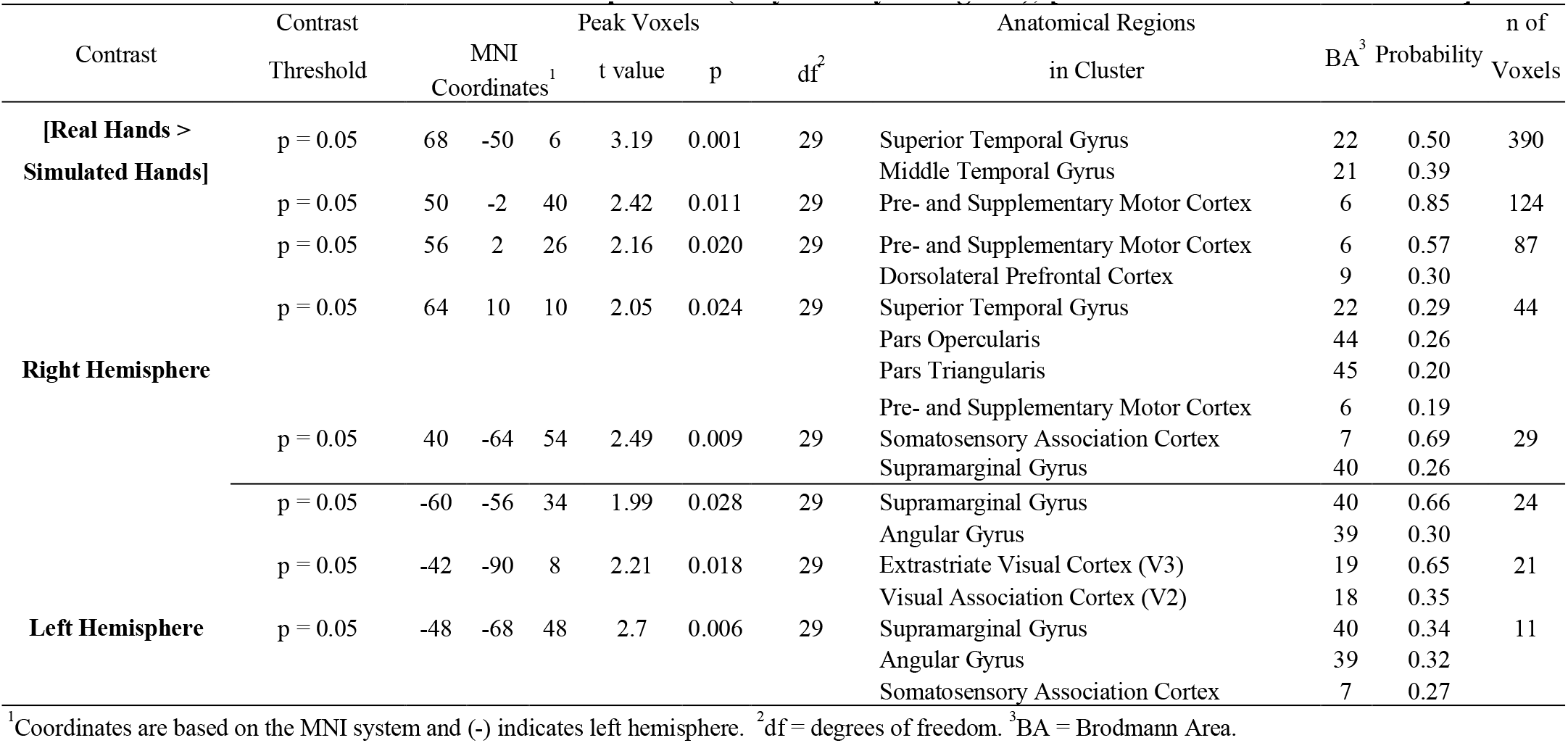
Voxel-wise GLM Contrast comparisons (Oxy+deOxyHb signals), [Real Hands > Simulated Hands].

### 3.2 Cross-Brain Neural Coherence

In addition to a cognitive social context, such as the theory-of-mind conventionally investigated with single participant paradigms (Carter & Huettel, 2013), it has been shown that the rTPJ also codes live interactions such as face-to-face processes (Kelley et al., 2021; Noah et al., 2020). Domain general processes for live interactions predicts cross-brain coherence between TPJ regions for social touch (Hasson et al., 2012). In accordance with this prediction, coherence was greater during the interactive real-hand tasks than simulated hand tasks (Figure 4).

**Figure 4.**
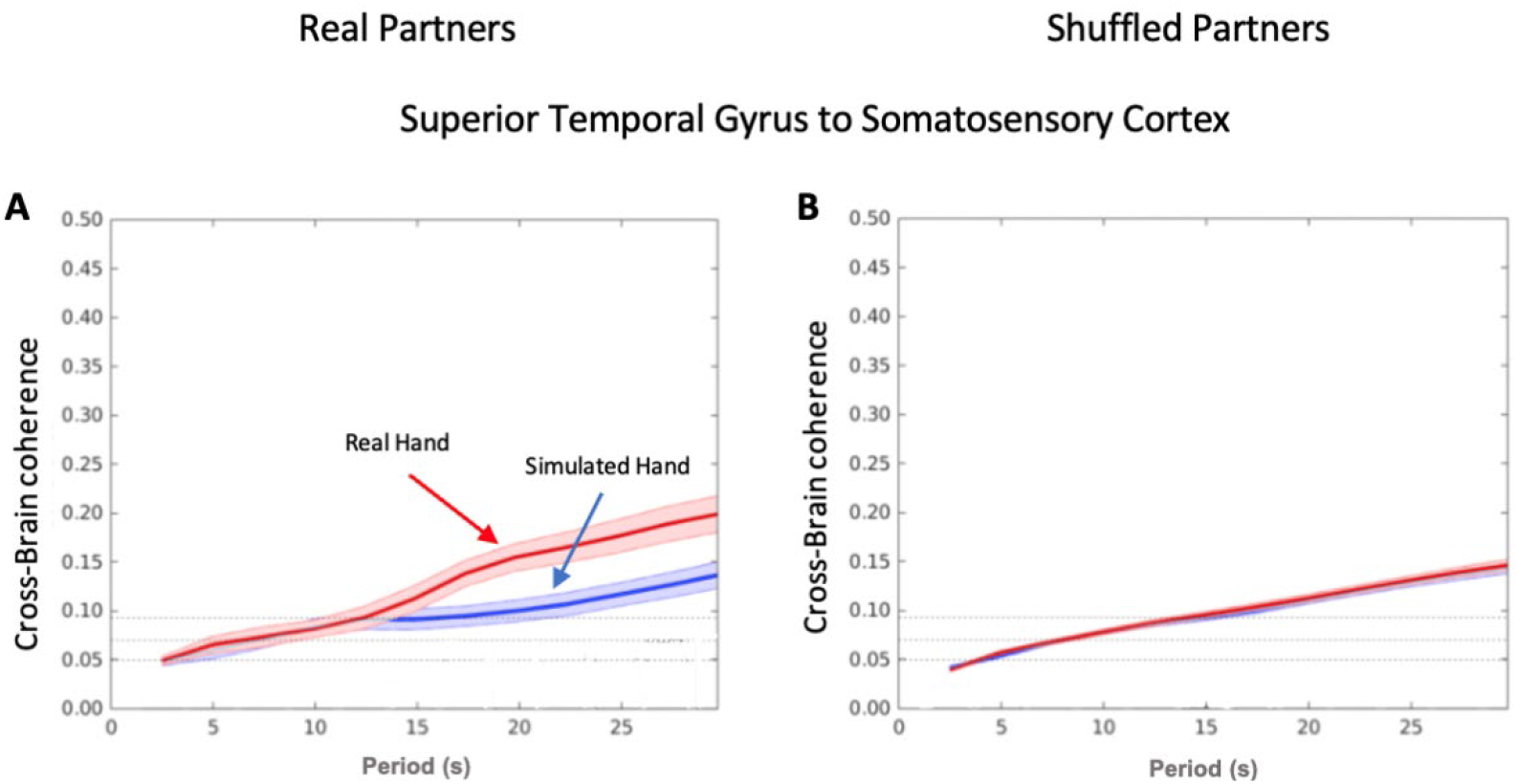
Brain-to-brain coherence between dyads (y-axis) is plotted against the temporal period (wavelength) (x-axis) for the real hand condition (right and left) (red) and the simulated hand condition (right and left) (blue) (surrounding colors ±1 SEM). A. depicts coherence between real partners whereas B. shows coherence between shuffled partners. Cross-brain coherence is highest in the real hand conditions between superior temporal gyrus to somatosensory cortex.

Temporal oscillations of hemodynamic signals decomposed into wavelet components (x-axis) are shown against the correlation between the signals of partners (y-axis) acquired during the joint tasks of grasping the real and the simulated hands, red and blue respectively (methods described previously (Hirsch et al., 2018; Hirsch et al., 2017; Noah et al., 2020). Cross-brain coherence between the superior temporal and somatosensory gyri was greater in the real hand task (red) in comparison to the simulated hand task (blue). Interestingly, this was true only for the case of real partners (Fig 4A), and not for shuffled partners (Fig 4B), (i.e., randomly paired with every other subject except the original partner) where no difference in coherence was found. This comparison is consistent with the interpretation that cross-brain signal coherence was due to the specific interactions between the participants performing the real hand clasp.

## 4 Conclusion

These findings advance our understanding of the neural systems that underlie social cognition and social touch consistent with a domain general function for the rTPJ. Although the rTPJ has been shown to play a role in social cognition, such as theory of mind (Carter & Huettel, 2013), as well as live face-to-face gaze (Kelley et al., 2021; Noah et al., 2020), this is the first documentation of a role in processing social touch and provides evidence in support of rTPJ as a domain general region for social and interactive functions. Further, findings of increased cross-brain coherence of rTPJ signals between partners during real hand clasping relative to simulated hand clasping are consistent with neural coupling between the interactive systems.

## 6 Author Contributions

J.H. designed and supervised this experiment. C.D., A.N, X.Z., J.L.C., and T.B. performed the experiment and analyzed the data. T.B. and J.H. wrote the manuscript.

## 7 Funding and Acknowledgements

This research was partially supported by the National Institute of Mental Health of the National Institutes of Health under award numbers 1R01MH111629 (PIs JH and JM); R01MH107513 (PI JH); 1R01MH119430 (PI JH). The content is solely the responsibility of the authors and does not necessarily represent the official views of the National Institutes of Health. All data reported in this paper are available upon request from the corresponding author.

## 8 Data Availability Statement

The datasets analyzed for this study will be made available upon request at fmri.org

