## Supplemental Table 1 for "Domain General Processes for Interactive Touch"

### Demographics for individual participants and dyads

| ID | Age | Female | Male | Right-handed | Asian | White | Latina | Biracial | Dyad Race |
|---|---|---|---|---|---|---|---|---|---|
| 1A | 27 | 1 | 0 | Yes | 0 | 1 | 0 | 0 | White & |
| 1B | 23 | 1 | 0 | Yes | 0 | 1 | 0 | 0 | White |
| 2A | 26 | 1 | 0 | Yes | 0 | 1 | 0 | 0 | Biracial |
| 2B | 41 | 1 | 0 | Yes | 0 | 0 | 0 | 1 | & White |
| 3A | 19 | 0 | 1 | Yes | 0 | 1 | 0 | 0 | Latina |
| 3B | 22 | 1 | 0 | Yes | 0 | 0 | 1 | 0 | & White |
| 4A | 34 | 1 | 0 | Yes | 0 | 1 | 0 | 0 | White & |
| 4B | 19 | 1 | 0 | Yes | 0 | 1 | 0 | 0 | White |
| 5A | 26 | 0 | 1 | Yes | 0 | 1 | 0 | 0 | Asian & |
| 5B | 20 | 0 | 1 | Yes | 1 | 0 | 0 | 0 | White |
| 6A | 26 | 1 | 0 | Yes | 0 | 1 | 0 | 0 | White & |
| 6B | 58 | 0 | 1 | Yes | 0 | 1 | 0 | 0 | White |
| 7A | 21 | 1 | 0 | Yes | 0 | 0 | 0 | 1 | Biracial |
| 7B | 20 | 1 | 0 | Yes | 0 | 1 | 0 | 0 | & White |
| 8A | 21 | 1 | 0 | Yes | 0 | 1 | 0 | 0 | White & |
| 8B | 32 | 0 | 1 | Yes | 0 | 1 | 0 | 0 | White |
| 9A | 18 | 1 | 0 | Yes | 1 | 0 | 0 | 0 | Asian & |
| 9B | 26 | 1 | 0 | Yes | 1 | 0 | 0 | 0 | Asian |
| 10A | 63 | 0 | 1 | Yes | 0 | 1 | 0 | 0 | Asian & |
| 10B | 21 | 1 | 0 | Yes | 1 | 0 | 0 | 0 | White |
| 11A | 20 | 0 | 1 | Yes | 0 | 1 | 0 | 0 | Asian & |
| 11B | 27 | 1 | 0 | Yes | 1 | 0 | 0 | 0 | White |
| 12A | 21 | 0 | 1 | Yes | 0 | 1 | 0 | 0 | White & |
| 12B | 29 | 1 | 0 | No | 0 | 1 | 0 | 0 | White |
| 13A | 57 | 0 | 1 | Yes | 0 | 1 | 0 | 0 | White & |
| 13B | 25 | 0 | 1 | Yes | 0 | 1 | 0 | 0 | White |
| 14A | 29 | 1 | 0 | Yes | 0 | 1 | 0 | 0 | White & |
| 14B | 46 | 0 | 1 | Yes | 0 | 1 | 0 | 0 | White |
| 15A | 20 | 1 | 0 | Yes | 0 | 1 | 0 | 0 | Asian & |
| 15B | 26 | 1 | 0 | Yes | 1 | 0 | 0 | 0 | White |
| **Age** | (years) | 19F | 11M | 29 Right | 6 | 21 | 1 | 2 | |
| Mean | 28.8 | | | 1 Left | | | | | |
| SD | ±12.2 | | | | | | | | |
| Median | 26 | | | | | | | | |
| Max | 63 | | | | | | | | |
| Min | 18 | | | | | | | | |

**Table S1.** Self-reported demographic information, including participant age; gender; handedness; race; and racial composition of dyads.
